# CAMO: A molecular congruence analysis framework for evaluating model organisms

**DOI:** 10.1101/2021.11.21.469371

**Authors:** Wei Zong, Tanbin Rahman, Li Zhu, Xiangrui Zeng, Yingjin Zhang, Jian Zou, Song Liu, Zhao Ren, Jingyi Jessica Li, Steffi Osterreich, Tianzhou Ma, George C. Tseng

## Abstract

CAMO provides a rigorous and user-friendly solution for quantification and mechanistic exploration of omics congruence in model organisms and humans. It performs threshold-free differential analysis, quantitative concordance/discordance scoring, pathway-centric investigation, and topological subnetwork detection. Instead of dichotomous claims of “poorly” or “greatly” mimicking humans, CAMO facilitates discovery and visualization of specific molecular mechanisms that are best or least mimicked, providing foundations for hypothesis generation and subsequent translational investigations.

---

As human studies often encounter numerous recruitment and ethical constraints, model organisms have played an indispensable role in pre-clinical research to understand pathogenesis and treatment response at the behavioral, cellular, and molecular levels. Their clinical validity and translational values are, however, long been debated with controversial opinions^1-4^. A notable example is the contradictory conclusions from two articles analyzing an identical transcriptomic response dataset in human and mouse inflammation^5,6^, with the former concluding “poorly mimicking” of the mouse model and the latter reporting “greatly mimicking”, which triggered further debates^7–9^. Although efforts have been made to compare or predict model organism responses using association analysis^5,6^, machine learning^10,11^, pathway enrichment^12,13^, or meta-analysis^14^ approaches, methods for exploring mechanistic insights are lacking. To meet the gap, we develop a <u>C</u>ongruence <u>A</u>nalysis of <u>M</u>odel <u>O</u>rganisms (CAMO) framework to evaluate omics congruence of animal models and aid mechanistic understanding, hypothesis generation, and translational guidance.

Fig. 1a-b overview CAMO’s pipeline. “Bayesian differential analysis” contrasting case/control or treated/non-treated groups are performed in human and mouse cohorts separately and “concordance and discordance scores” (abbreviated as c-score and d-score) are calculated (Fig. 1a), reflecting degree of cross-species congruence and discrepancy. The threshold-free Bayesian differential model transforms p-values obtained from routine pipelines, such as LIMMA or DEseq2, into differential posterior probabilities, which in turn are input to cross-species c-score/d-score calculation based on a stochastic version of confusion matrix and F-measure in the machine learning setting with p-values assessed by permutation. When multiple cohorts are jointly analyzed, c-scores and d-scores are calculated for all pair-wise studies in each pathway. Next, the “Mechanistic investigation and hypothesis generation” component can perform “pathway knowledge retrieval” and “topological gene module detection” (Fig. 1b). In pathway knowledge retrieval, top (concordance or discordance) enriched pathways are clustered with similar congruence patterns across studies and a text mining algorithm is implemented to retrieve representative keywords to interpret each pathway cluster. In each pathway, a community detection algorithm can further identify concordant or discordant subnetworks based on topological regulatory information.

**Figure 1.**
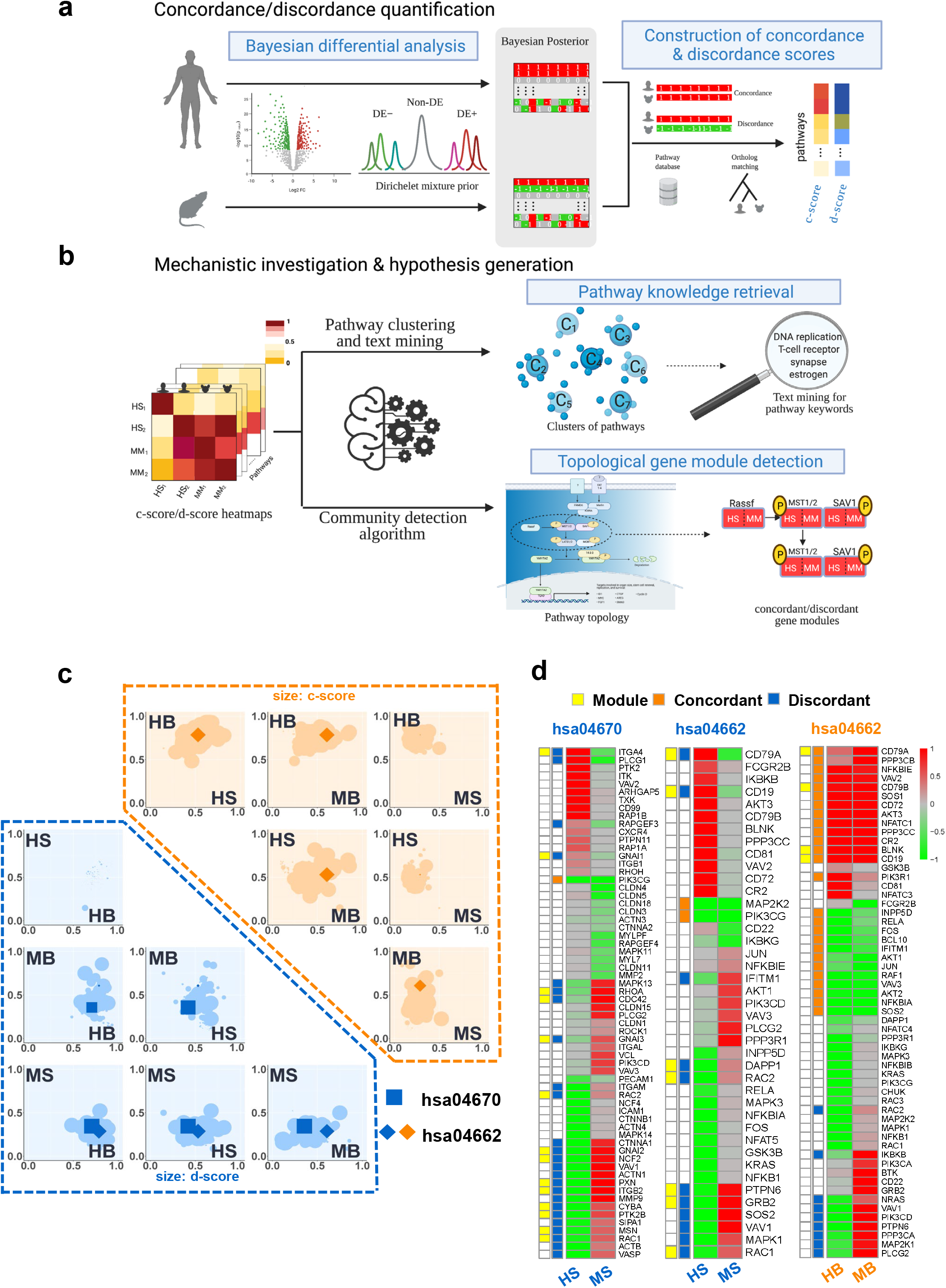
Workflow of the “CAMO” framework with application results from Case Study 1. (a) Procedures to calculate genome-wide and pathway level c-scores and d-scores for a pair of human study (HS) and mouse model (MM). (b) Downstream machine learning and bioinformatics interactive visualization tools for pathway knowledge retrieval and topological gene nodule detection. (c) Summary of DE evidence with pathway level c-scores (orange in the upper right region) and d-scores (blue in lower left region). X- and Y-axes represent the average DE posterior probabilities, and size of dots represents the magnitudes of c-scores (orange) or d-scores (blue). Two example pathways are highlighted using different shapes (“◇”: hsa04662 - KEGG: B cell receptor signaling pathway; “☐”: hsa04670 - KEGG: Leukocyte transendothelial migration). (d) Gene-wise heatmap of posterior mean of DE indicators of the HS-MS comparison in hsa04670, HS-MS in hsa04662 and HB-MB in hsa04662. Genes identified by community detection algorithm (yellow) and genes with concordant (orange) or discordant (blue) are shown in two columns beside the heatmaps.

Our first case study re-evaluates the contradicting papers in human-mouse transcriptomic response to inflammation. Supplementary Table 1 lists 12 studies in human and mouse (Burns, Infection, Trauma, Sepsis, LPS and ARDS), abbreviated as HB, HI, HT, HS, HL and HA, and MB, MI, MT, MS, ML and MA (H for human and M for mouse). Supplementary Table 2 summarizes analytical differences and arbitrary thresholds in the two papers that may have contributed to the contradicting conclusions. CAMO avoids such subjective analytical thresholds and decisions and extends the investigation into pathways and gene regulatory modules for insightful mechanistic understanding. Supplementary Table 3 contains genome-wide c-scores and d-scores of pair-wise studies and the c-score-based multidimensional scaling (MDS) plot in Supplementary Fig. 1 shows that four human studies HB, HI, HT and HS resemble each other well (c-scores=0.25~0.52). MI, MB and MT are relatively similar to the four human studies while MA, ML and MS have almost no genome-wide congruence to human, implying that cross-species congruence is condition specific. Unlike most of the human studies, the six mouse studies generally do not mimic each other, implying complexity and high variability of mouse models in inflammatory diseases. We next apply consensus tight clustering to 219 concordance enriched pathways and identify four pathway clusters (Supplementary Fig. 2-3). For example, the heatmap and text mining results show high congruence between human (HB, HS, HT and HI) and mouse models (MB, MI and MT) in both innate and adaptive (e.g., B and T cell related) immunity (Supplementary Fig. 3 and Supplementary Table 4). Despite the difference in neutrophil and lymphocyte abundance, it has been reported that the overall immune system is relatively similar in mouse and human^15^. Cluster IV shows mouse models do not mimic human studies in ribosome and protein translation. Such findings agree with earlier studies that profound cross-species differences exist in translation machinery^16^.

CAMO next provides interactive exploration in the shiny app to select pathways and zoom in for regulatory topological visualization. Fig. 1c shows differential expression (DE) evidence and c-scores/d-scores of individual pathways in pair-wise comparisons of HB, HS, MB and MS. Fig. 1d displays gene-specific concordance or discordance information in two selected KEGG pathways. Fig. 2a highlights the KEGG gene-gene regulatory topological plot for “Leukocyte transendothelial migration” pathway (hsa04670) with side-by-side display of the differential regulation signals (red for up-regulation and green for down-regulation) in HS and MS. The community detection algorithm identifies a module of 14 DE genes (RHOA, PTK2B, RAC2, RAC1, CDC42, ITGA4, ITGB2, MSN, PXN, NCF2, CYBA, GNAI1, GNAI2 and GNAI3) with opposite effect sizes (green in HS and red in MS or vice versa; p = 0.002). The co-localized discordant module is directly related to cell motility and direct sensing, a critical function that allows leukocytes to attach to the vessel wall to initiate immune response during inflammation^17^. The striking mouse-human discordant result may reflect the discrepancy in proportions of different cell types of blood leukocytes between human and mouse as pointed out in a previous critique^9^. The topological plot for “B cell receptor signaling pathway” (hsa04662) (Fig. 2b) shows a gene module of 7 discordant genes (PTPN6, DAPP1, CD79A, RAC1, RAC2, GRB2, CD19; p=0.009). CD79A and CD19 are antigen receptors on B cell membrane to regulate signaling molecules, such as GRB2 and RAC family, with important roles in the regulation of cell growth and movement. On the other hand, Fig. 2c shows generally concordant DE signals between HB and MB (both red or both green) in the B cell membranes receptor and signaling, including CD72, CD79A, CD79B, IFITM1, CD19, CR2 and BLNK. Previous literature has pointed out the similarity but also significant differences between mouse and human immunology, specifically in B cell development^15^.

**Figure 2.**
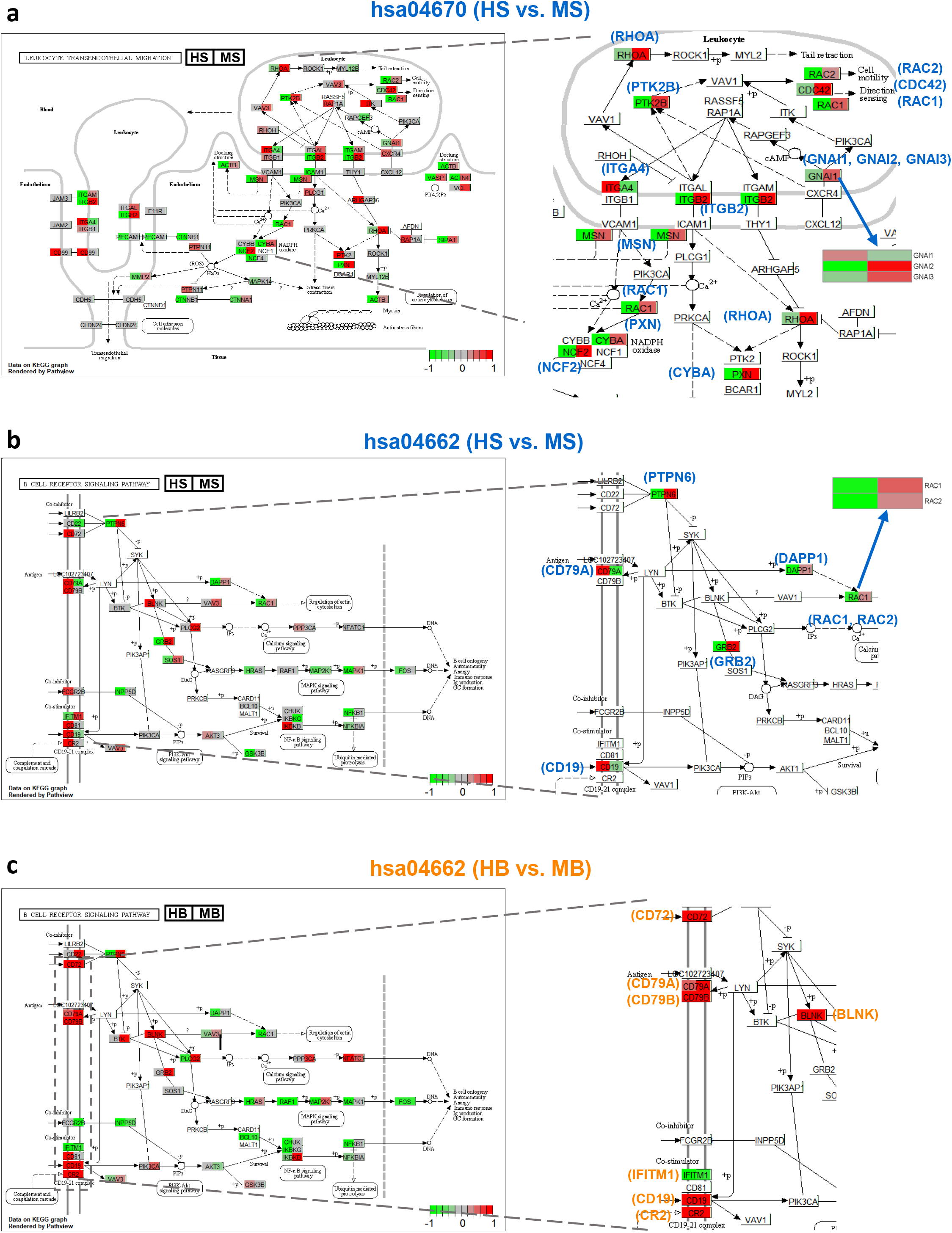
Pathway topology plots of the selective pathways Case Study 1. (a). hsa04670 (HS-MS), (b). hsa04662 (HS-MS) and (c). hsa04662 (HB-MB). Pop-out plots represent the co-localized concordant/discordant modules identified from the pathway topology by the community detection algorithm. Colors in the nodes refer to the posterior mean of DE indicators in each corresponding study pair (red for up-regulation and green for down-regulation).

The second case study evaluates transcriptomic congruence of *C. elegans* (ce) and *D. melanogaster* (dm) in developmental stages using the modENCODE data^18,19^, where five developmental stages in each species are confirmed by hierarchical clustering (Supplementary Fig. 4): early embryo (ce.e0), mid embryo (ce.e1), late embryo (ce.e2), larvae (ce.lar), dauer (ce.dau) using *C. elegans* adult as the reference; early embryo (dm.e0), mid embryo (dm.e1), late embryo (dm.e2), larvae (dm.lar), pupae (dm.pup) using female Drosophila adult as reference. Supplementary Fig. 5a shows MDS plot of genome-wide c-scores for the five Drosophila and five *C. elegans* stages. The y-axis shows a clear separation between the two species. The x-axis presents a developmental transition in the embryonic stages e0→e1→e2, while the larvae and pupae/dauer stages are not exactly ordered. Adjacent developmental stages are found to be more similar within species. ce.e2 shows some resemblance with all developmental stages in Drosophila except for dm.e0. This unintuitive result is better visualized by an intriguing bipartite graph between Drosophila and *C. elegans* stages (Supplementary Fig.5b) by creating solid edges when pair-wise genome-wide c-scores are greater than 0.1 (Supplementary Table 5). We first observe reasonable within-stage cross-species resemblance (i.e., solid yellow edges: ce.e0—dm.e0, ce.e1—dm.e1, ce.e2—dm.e2, and ce.e2—dm.e1; dashed yellow edge: ce.lar—dm.lar) and then identify surprising cross-stage resemblance between species (i.e., purple edges: ce.dau—dm.e2, ce.e2—dm.lar and ce.e2—dm.pup). Resemblance of ce.dau—dm.e2 has been suggested by the original modENCODE paper^18,19^. Resemblance of ce.e2—dm.lar and ce.e2—dm.pup confirms the second large wave of cell proliferation and differentiation in Drosophila’s life cycle.

From 269 concordance enriched pathways, consensus tight clustering identifies six pathway clusters (Supplementary Fig. 6,7), including Cluster III related to cell cycle and DNA replication with cross-species late-stage congruence, Cluster IV related to estrogen and hormone in *C. elegans*, and Cluster II specific to Drosophila developmental stages. Supplementary Fig. 8 shows DE evidence and c-scores/d-scores in ce.e2, ce.dau, dm.e2 and dm.pup. Pathways “Homologous recombination” (KEGG: cel03440) and “Mismatch repair” (KEGG: cel03430) exhibited high concordance between ce.e2 and dm.e2 (Supplementary Fig. 9), implying similar molecular events taking place in late embryo stage for both species. The pathway “Nucleotide-binding domain, leucine rich repeat containing receptor (NLR) signaling pathways” (Reactome: R-CEL-168643) exhibits discordance between ce.dau and dm.pup (Supplementary Fig. 9). The NOD1/2 and inflammasomes components of the pathway are both related to the innate immune system, the first line of defense against invading microorganisms that are present in the pupae stage of Drosophila but not in the dauer stage of *C. elegans*^20^.

Notwithstanding the fact that human and animal studies are both fundamental in disease and drug investigation, objective and interactive congruence evaluation to distinguish and integrate cross-species omics information is currently lacking. We expect that CAMO and its future extension to multi-omics and single cell data will provide profound cross-species mechanistic understanding to improve animal models and to accelerate treatment development in human diseases.

## ONLINE METHODS

### Threshold-free Bayesian differential analysis

CAMO applies a Bayesian mixture (BayesP) model to derive differential expression posterior probabilities and to facilitate calculation of c-scores and d-scores in the next section, where the input of BayesP can be DE results from any conventional pipeline (e.g., “limma” for microarray and “DESeq2” for RNA-seq are used in this paper). Specifically, the one-sided p-values, *p_g_* for gene *g*, from conventional DE analysis are first transformed to z-scores *Z_g_* = Φ^−1^(*p_g_*) (Φ^−1^(·) is inverse CDF of standard normal distribution), which incorporates information of both statistical significance and directionality. BayesP adopts the following three-component Gaussian mixture model for up-regulated, down-regulated and no-change genes similar to Huo et al. 2019 ^21^:

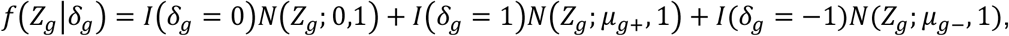

where *δ_g_* is the DE indicator of gene *g* (*δ_g_* = 1 indicates up-regulation, *δ_g_* = −1 for down-regulation and *δ_g_* = 0 for no change), and *μ_g+_* and *μ_g-_* are the grand means of z-scores of the up-regulated and down-regulated groups. We assume a non-parametric Dirichlet process prior on the grand means: 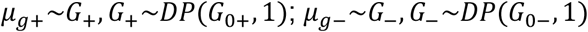, where *G*_0+_ (*G*_0-_) denotes a left (right) truncated *N*(0,10^2^) and 1 is the concentration parameter of the Dirichlet process. A Chinese Restaurant Process is used to update *δ_g_*, where we define an auxiliary component variable *C_g_* ∈ {…,-2,-1,0,1,2,…} such that *C_g_* = 0 indicates *δ_g_* = 0, *C_g_* > 0 indicates *δ_g_* = 1 and *C_g_* < 0 indicates *δ_g_* = −1. The prior for *δ_g_* is specified as: *P*(*δ_g_* ≠ 0) = *π_g_,P*(*δ_g_* = 1|*δ* ≠ 0) = *ρ_g_*; 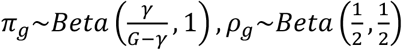, where 1 -*π_g_*, *π_g_ρ_g_* and *π_g_*(1 - *ρ_g_*) are the prior probabilities of being no-change, up-regulated and down-regulated genes respectively.

Markov chain Monte Carlo (MCMC) using Gibbs sampling is used to update all parameters (*π_g_, ρ_g_, δ_g_*) sequentially as follows:

1. Update 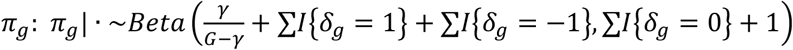
2. Update 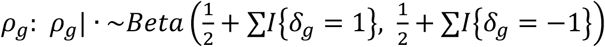
3. Update 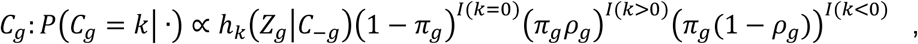 where *h_k_*(*Z_g_|C_-g_*) is derived according to Neal 2000 ^22^.
4. Update *δ_g_*: *δ_g_* = *sgn*(*C_g_*) where *sgn*(·) is the sign function.

The empirical distribution of the posterior probabilities of *δ_g_* will be used to derive the cross-species c-scores and d-scores later.

### Deterministic version of cross-species c-scores and d-scores

The foundation of cross-species c-scores and d-scores comes from a natural definition of confusion matrix and F-measure in machine learning (Supplementary Table 7) when human and mouse DE status of up-regulation (Ω^*H*+^ and Ω^*M*+^), down-regulation (Ω^*H-*^ and Ω*^M-^*) and no change (Ω^*H*0^ and Ω^*M*0^) are deterministically known, where 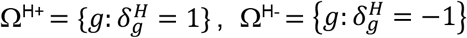 and 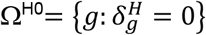 in human, and similarly for mouse. Denote by *a, e* and *i* the number of cross-species DE concordant genes: *a* = #(Ω^*H*+^ ⋂ Ω^*M*+^) (number of concordant up-regulated genes), *e* = #(Ω^*H*0^ ⋂Ω^*M*0^) (number of concordant no-change genes), and *i* = #(Ω^*H*-^ ⋂ Ω^*M*-^) (number of concordant down-regulated genes). The numbers of DE discordant genes can be similarly defined for *b, c, d, f, g, h* in the contingency table. From the viewpoint of machine learning prediction benchmark assuming we use mouse DE status to predict human DE status, one can define concordance sensitivity_C_ (a.k.a. recall_C_)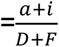, and 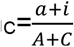; when we focus on cross-species concordant DE genes, where *A* = #(Ω^*M*+^), *C* = #(Ω^*M*-^), *D* = #(Ω^*H*+^) and *F* = #(Ω^*H*-^). In sensitivity_C_, we calculate the number of concordant DE genes (i.e., *a* + *i*) among the true human DE genes (i.e., *D* + *F*). Similarly, precisionC is defined as the number of concordant DE genes (i.e., *a* + *i*) among the claimed mouse DE genes (i.e., *A* + *C*). We define the raw DE concordance score between human and mouse as the F-measure: *c*’ =2(precision_C_ x recall_C_)/(precision_C_+recall_C_). Similarly, we can focus on DE discordant genes (i.e., genes up-regulated in human but down-regulated in mouse or vice versa) and define 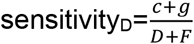; and 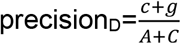. The raw DE discordance score between human and mouse becomes: *d*’ =2(precision_D_ x recall_D_)/(precision_D_+recall_D_). In addition to F-measure, we can also use Youden index (=sensitivity+specificity-1) or the geometric mean of sensitivity and specificity, where 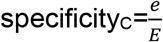 and 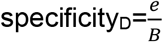. When there is no reference study specified or under the general multi-cohort scenario, the F-measure is a better choice among the three because it is symmetric no matter which species is taken as the reference. With simple algebraic calculation, one can show that 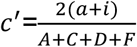 and 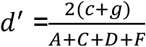. Similar to Rand index used to evaluate clustering similarity and the adjusted Rand index subsequently developed ^23^, although both *c*’-score and *d*’-score range between 0 and 1, their expected value under null hypothesis (i.e., no resemblance between mouse and human) is not 0, making the interpretation difficult. To account for this pitfall, we adjust the scores to have maximum value at 1 for perfect resemblance and expected value at 0 when no resemblance exists using a linear transformation: 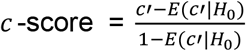 and 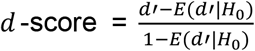, where *H*_0_ is the null hypothesis when mouse and human have no resemblance, 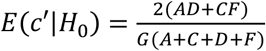 and 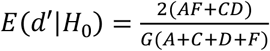 by computing the expected counts from the table margins for each cell (e.g. 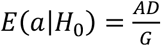).

### Data-driven estimation version of c-scores and d-scores

In practice, the underlying true DE statuses (Ω^*H*+^, Ω^*H*0^, Ω^*H*-^) and (Ω^*M*+^, Ω^*M*0^, Ω^*M*-^) are not known and are inferred from data. As previously mentioned, cross-species congruence analysis by applying arbitrary p-value/FDR and fold change cutoffs can lead to subjective bias and inconsistent conclusions ^5,6^. In CAMO, we infer Bayesian posterior probabilities and plug into the deterministic definition of c-scores and d-scores. Specifically, denote by 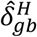 the simulated estimation of 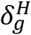 in the *b*-th MCMC iteration for gene *g* in the human study and similarly 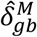 for the mouse study. The unbiased estimators are obtained as: 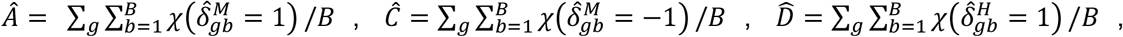 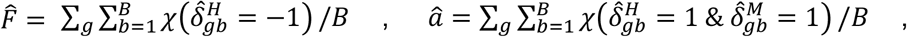 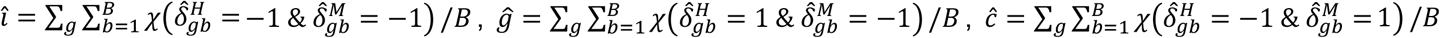, where *B* is the number of (post burn-in) MCMC simulations and *χ*(·) is the indicator function taking value 1 if the statement is true and 0 otherwise. c-score and d-score are estimated by plugging these estimators into their deterministic definitions.

### Pathway-specific c-scores and d-scores

The aforementioned c-score and d-score estimations are calculated in the genome-wide scale. Since the cross-species congruence can vary by biological pathways, we analogously define pathway-specific c-scores and d-scores by constraining the calculation to each pathway. One major modification is when calculating the expected raw score under null hypothesis, a subsampled (sample without replacement) gene set with equivalent size of the target pathway is used to calculate 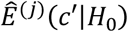 and 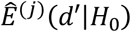 in the *j*-th sampling. We then estimate 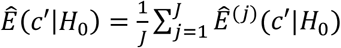 and 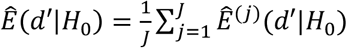 to better represent the genome-wide status.

### Statistical significance (p-value) assessment of c-score and d-score

We assess p-values of genome-wide and pathway-specific c-scores and d-scores by permutation analysis. Specifically, we randomly permute cross-species ortholog gene annotation, so no cross-species congruence exists under the null hypothesis and the procedure is repeated for *T* times. The p-values are calculated as 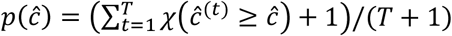 and 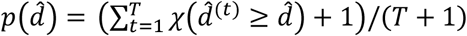, where 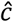 and 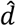 are the calculated c-score and d-score, and 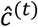 and 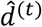 are the derived c-score and d-score in the *t*-th permutation. Note that we count 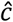 and 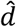 as one of the permutation observations to avoid obtaining zero p-values ^24^. Benjamini-Hochberg (BH) procedure ^25^ is applied to adjust for multiple comparisons of testing many pathways. Both pathway specific c-scores and d-scores and their associated p-values are essential in CAMO to identify pathways most or least mimicked by the animal model and to investigate the underlying mechanism.

### Pathway clustering and text mining

In CAMO, congruence analysis is evaluated in a pair of studies. When we assess M studies, CAMO will create 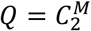 congruence analysis results. Depending on selection of pathway databases, hundreds or up to thousands of pathways are assessed for c-scores and d-scores, and the result can contain high redundancy since different pathway databases may describe a related biological function using similar gene sets. Denote by *C*_*K*×*Q*_ = {*c_kq_*} and Θ_*K*×*Q*_ = {*θ_kq_* = -*log*_10_*p*(*c_kq_*)} the matrices of c-scores and associated minus-log-transformed p-values of the *Q* congruence comparisons in *K* pathways. Note that large value of *θ_kq_* represents high concordance in the *q*-th congruence evaluation of pathway *k*. To further decipher and interpret pathway-specific congruence result, We consider dissimilarity (Euclidean distance 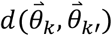) between 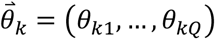 and 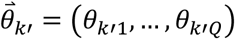 of pathways *k* and *k*’ and cluster the statistically significant pathways (i.e., meta-analyzed q-values by Fisher’s method across *Q* comparisons *≤* 0.05) using a consensus tight clustering algorithm. The algorithm uses the resampling-based consensus clustering ^26^ for identifying stable patterns in data followed by removing the scattered pathways with low silhouette width ^27^ iteratively until all pathways’ silhouette widths are above a certain cutoff (e.g., 0.1) to improve the tightness of clusters. Pathways with similar concordance patterns across the *Q* pairwise comparisons of the *M* studies are clustered together to reduce redundancy and facilitate further investigation. A heatmap of the matrix Θ_*K*×*Q*_ sorted by pathway clusters is shown to visualize the concordance patterns in different clusters (Supplementary Fig.3a,7a). A multidimensional scaling (MDS) algorithm is applied to the dissimilarity matrix generated from Θ_*K*×*Q*_ for visualization (Supplementary Fig.3b,7b). Finally, the co-membership heatmaps are used to summarize the proportion of significantly concordant pathways within each pathway cluster between each pair of studies (Supplementary Fig.3c,7c).

We next apply an automated text mining pipeline to extract summary annotations and retrieve knowledge from each pathway cluster ^28^. The method first collects names and summary descriptions of all pathways and extract noun phrases after filtering of biologically redundant phrases and merging synonyms using R packages *spacyr, tm, textstem* and *wordnet*. The remaining noun phrases are tested for whether significantly enriched in selected pathway clusters by performing a permutation test on a cluster score weighted by length of pathway description. The output of text mining includes a list of key phrases most enriched and the corresponding permutation p-values for each pathway cluster.

### Individual pathway topology and co-localized concordant/discordant gene module detection

Pathway databases such as KEGG ^29^ and Reactome ^30^ provide pathway topological graphs to visualize involved genes, gene-gene interactions and regulatory information in the pathway. In the R-shiny interface of CAMO, we map and incorporate the gene-based concordance/discordance inference results in mouse-human comparison to the pathway graph to allow users for visual mechanistic investigation of the local concordance/discordance pattern. For pathways from KEGG, we use R package “Pathview” ^31^ to render the topology graph and integrate the concordance/discordance information. For pathways from Reactome, we developed our own tool to first retrieve and parse the pathway topology (.sbgn file) from Reactome database using the Python *minidom* parser (https://docs.python.org/3/library/xml.dom.minidom.html). Then, each node is colored by its posterior mean of DE assignment in the two studies side by side using the Python Imaging Library (https://pillow.readthedocs.io/en/stable/).

To avoid visual bias and to further investigate the local concordance/discordance pattern inside the pathway, we develop a community detection algorithm to identify closely connected concordant or discordant gene modules based on shortest path distance in the graph, where the unweighted graph is constructed using R packages “KEGGgraph” ^32^ and “xml2”, and the shortest path matrix is calculated by R package “igraph” ^33^. Exhaustive search algorithm is implemented to identify the concordant/discordant gene set with the smallest average shortest path at a given module size. However, for a pathway with a large number of concordant/discordant genes (e.g., size>30), exhaustive search is not feasible and a simulated annealing (SA) algorithm is used for fast search. We define the initial temperature *T*_0_, the temperature multiplier *μ*, the number of iterations for reaching equilibrium *N*, and the final temperature *T_f_*. Intuitively, *T*_0_ controls the acceptance of a trial assignment, *N* controls the maximum number of annealing iterations, *μ* controls speed of cooling down process for *T*_0_ to drop below *T_f_* and stop the process. The annealing process is harder when *T*_0_ is larger, *μ* is smaller and *N* is larger. Real data evaluation shows *T*_0_ = 10, *μ* = 0.95, *T_f_* = 1*e* - 5 and *N* = 1000 work well in general. The energy function is defined as the average shortest path of the current module denoted as *avgSP*(*G_m_*) where *m* is a given module size. A trial module 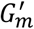 is proposed by randomly substituting one node in *G_m_* to another one in the searching space. If 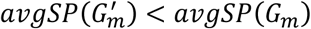, *G_m_* will be accepted immediately, otherwise it will initiate the annealing process. Another parameter *R* is introduced to control the total iterations in case it keeps hitting 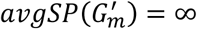 throughout the algorithm in a very sparse graph. Detailed algorithm can be found in the Supplementary notes 1. When it is applied to a spectrum of module sizes, to further improve the performance, at each module size *m*, the algorithm runs *x* times and the top *y* results are stored and passed to the next *m* +1 scenario as initials. Borrowing initial values from the previous step allows this procedure to converge faster and *y* >1 helps to robustize the procedure when multiple close-to-optimal solutions exist. The overall SA algorithm including the initialization procedure is summarized in the Supplementary notes 2.

Permutation test is performed to assess the p-value of identified concordant or discordant gene modules. Specifically, for an observed smallest *avgSP_m_* at module size *m*, gene modules of the same size are sampled without replacement from the searching space *B* times resulted in 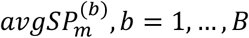. The p-value is derived as 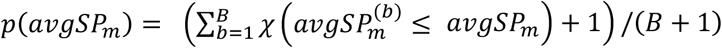 and the standard deviation of the *p*(*avgSP_m_*) is estimated as 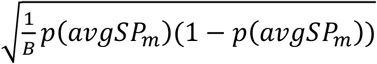 by regarding the permuted p-value as the mean estimate of *B* approximately independent Bernoulli trials.

In Case Study 1, we apply this local community detection algorithm to KEGG pathways hsa04670 and hsa04662 to identify discordant modules using exhaustive search. An elbow plot of (*avgSP_m_*) over *m* is generated from *m* = 4 to the cardinality of searching space i.e., the total number of discordant genes (Supplementary Fig. 10). The SA algorithm with *x* = 1 and *y* = 1 generates similar results as the exhaustive search. The maximum module size whose p-value is within 2 standard deviations of the minimum p-value is reported (12 nodes containing 14 genes in hsa04670 and 6 nodes containing 7 genes in hsa04662). Corresponding KEGG topology plots with highlighted gene modules are shown in Fig. 2. We recommend users to consider the p-value elbow plot, KEGG topology plots together with their biological insights in determining an appropriate module size to report.

### Data availability

The datasets used in Case Study 1^34^ are publicly available on the NCBI GEO database with accession numbers in Supplement Table 1. The datasets used in Case Study 2 are available at http://jsb.ucla.edu/software-and-data.

## Code availability

An open-source R package can be downloaded from https://github.com/weiiizong/CAMO.

## Acknowledgements

The authors acknowledge Diane Litman for helpful discussions. WZ, TR, LZ, YZ, JZ, ZR, TM and GCT are supported by R21LM012752 and R01CA190766. JJL is supported by R35GM140888 and R01GM120507.

## Author contributions

TM and GCT conceived the framework. WZ, TR, LZ, XZ, YZ, JZ, SL, ZR, TM and GCT developed the statistical and computational methods, and performed the analyses. WZ, JJL, SO, TM and GCT discussed the results and contributed to the final manuscript presentation. All authors proofread and agreed with the final manuscript.

