## Supplementary Figures and Tables for "CAMO: A molecular congruence analysis framework for evaluating model organisms"

**Supplementary Figure 1.** MDS plot based on genome-wide pairwise c-scores of the 12 studies in Case Study 1.

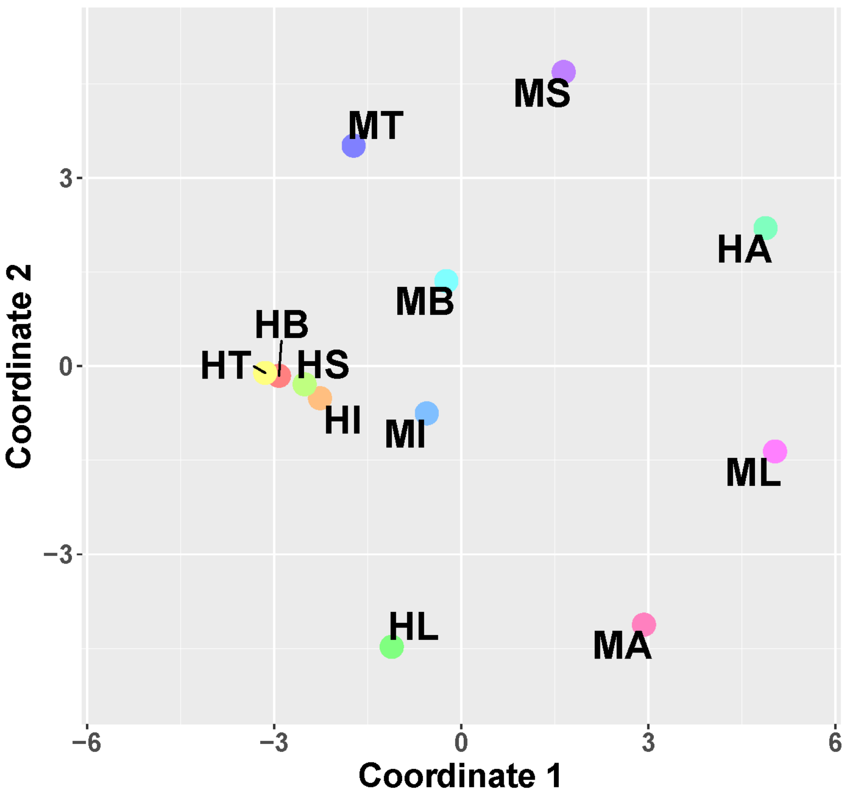

**Supplementary Figure 2.** Consensus clustering plots to determine the number of clusters in Case Study 1. (a). Consensus CDF plot. (b). Consensus clustering scree plot. The result clearly chooses to select four pathway clusters.

a

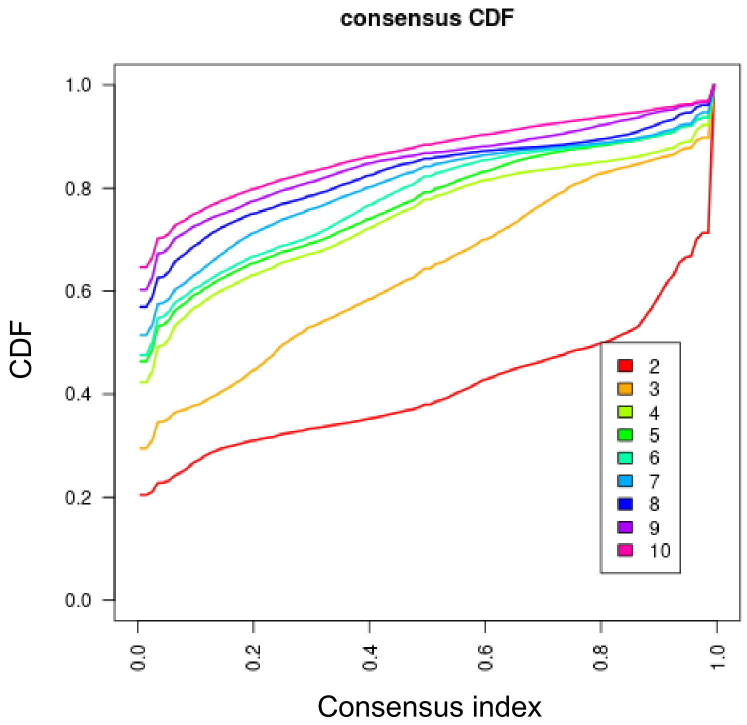


b

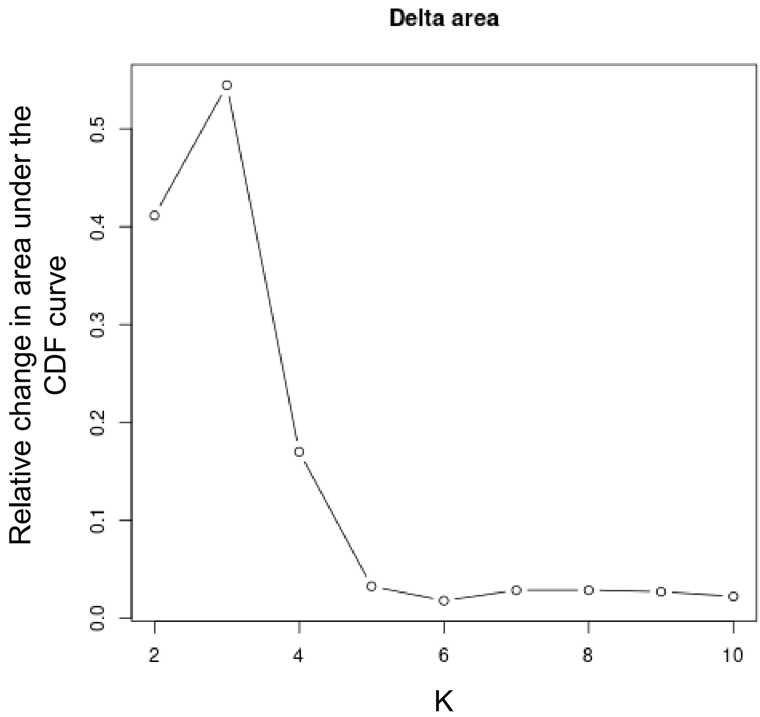

**Supplementary Figure 3.** Visualizations of pathway clusters in Case Study 1. (a). Heatmap of the –log10p(c-scores) matrix by red-green gradient color. Columns are pairs of studies evaluated for concordance and rows are pathways ordered by four clusters (red, green, light blue and purple) and the remaining singleton (un-clustered) pathways (gray). (b). MDS using distances based on the –log10p(c-scores) matrix of pathways colored by cluster membership. (c). Co-membership heatmaps of significantly concordant pathways within each pathway cluster between each pair of studies. Each cell is colored by the proportion of pathways in which the two corresponding studies are clustered together based on the pathway level c-scores. Each heatmap is annotated with key noun phrases from the text mining algorithm.

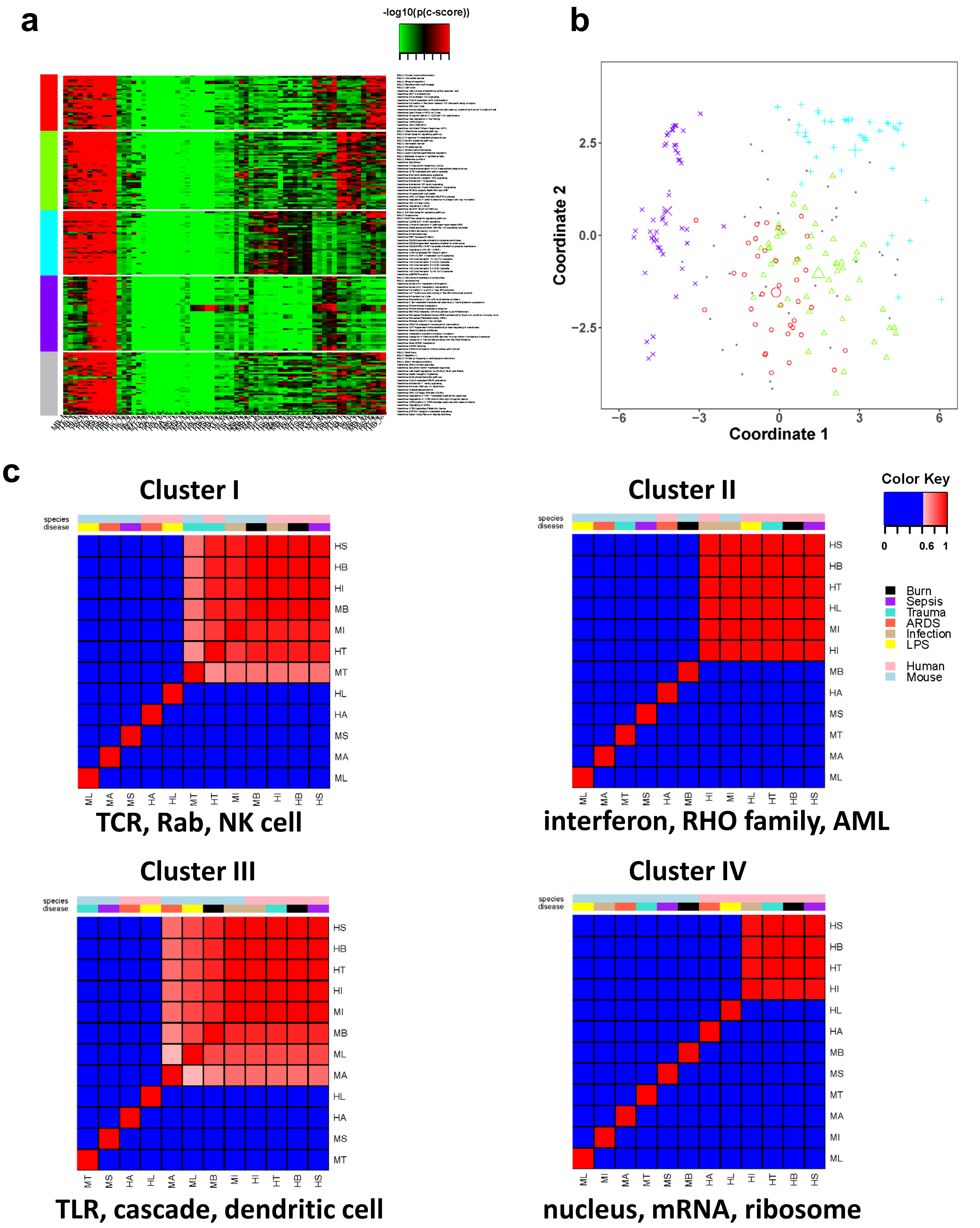

**Supplementary Figure 4.** Hierarchical clustering heatmap of (a) Caenorhabditis elegans' developmental stages and (b) Drosophila melanogaster's developmental stages. Embryo stage in both species breaks into three clearer sub-stages: early embryo, mid-embryo and late embryo. Female and male adult samples in Drosophila are separated. Female adults are used as base-line reference in this case.

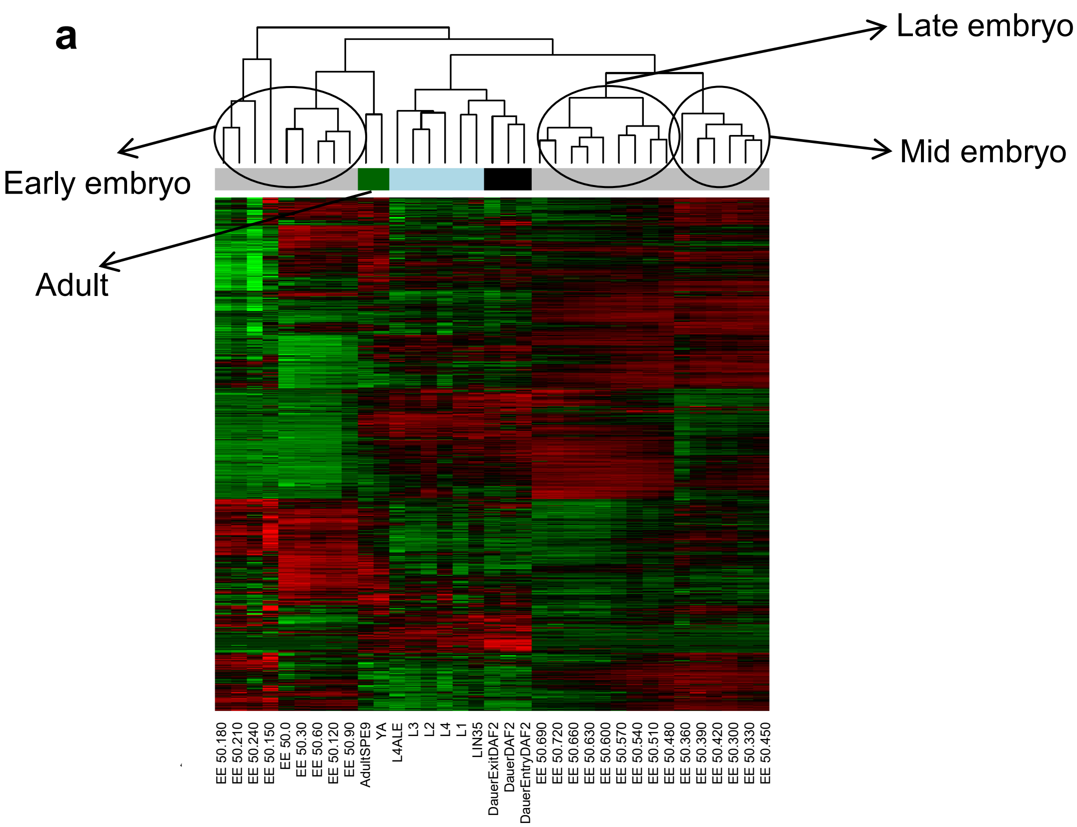


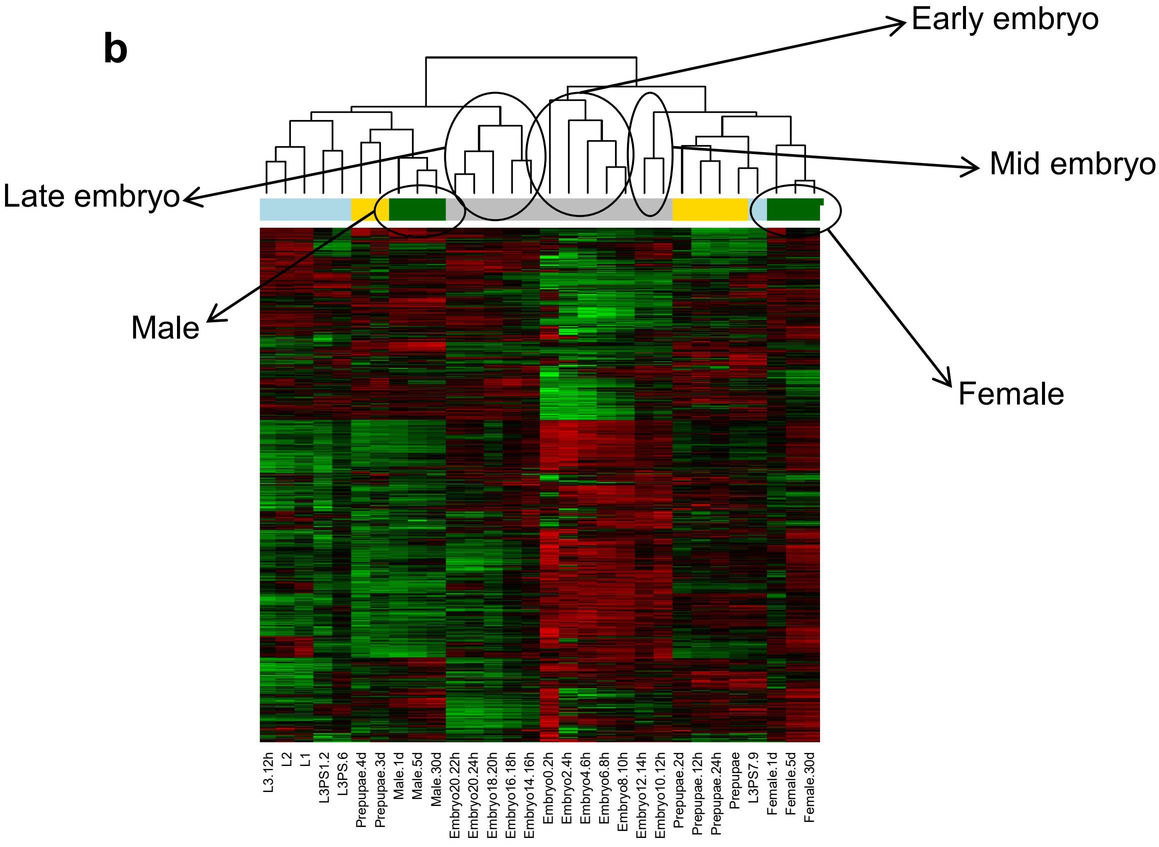

**Supplementary Figure 5.** (a). MDS plot based on genome-wide pairwise c-scores of the 10 studies in Case Study 2. (b). Bipartite graph between Drosophila and C. elegans where solid edges are draw when the genome-wide c-scores between any pair of cross-species stages are greater than 0.1. The yellow dashed line indicates a slightly weaker within-stage concordance with c-score=0.087.

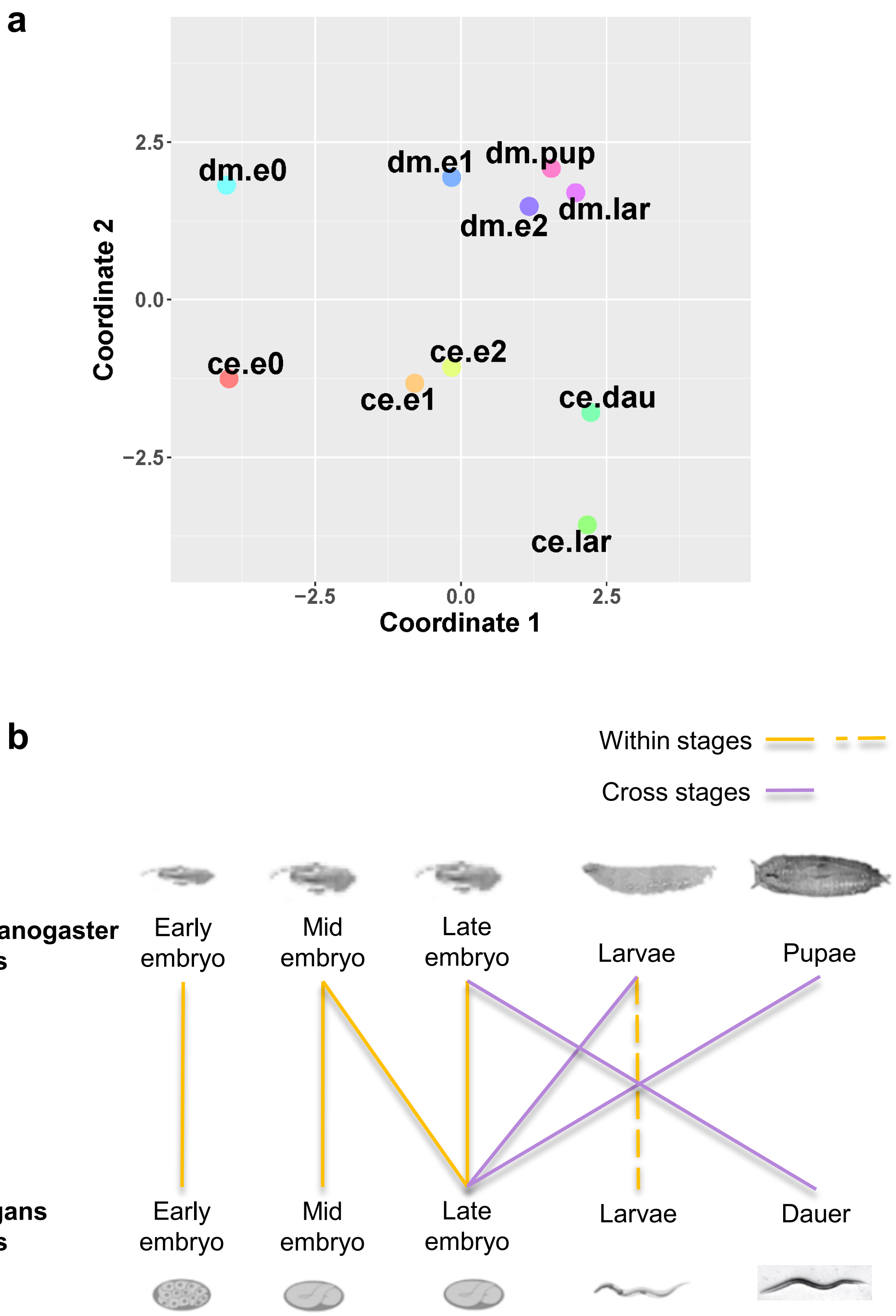

**Supplementary Figure 6.** Consensus clustering plots to determine the number of clusters in Case Study 2. (a). Consensus CDF plot. (b). Consensus clustering scree plot.

a

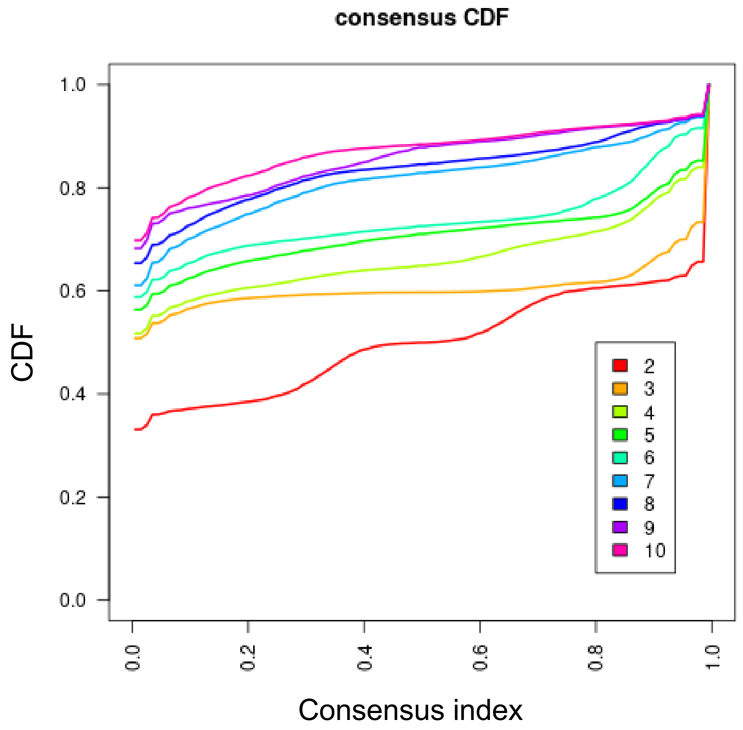


b

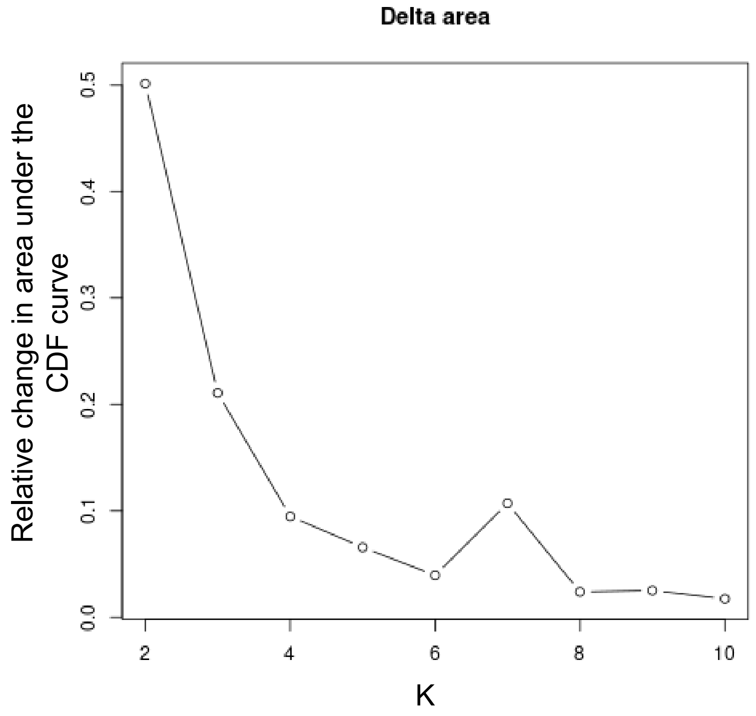

**Supplementary Figure 7.** Visualizations of pathway clusters in Case Study 2. (a). Heatmap of the –log10p(c-scores) matrix. Columns are pairwise studies and rows are pathways ordered by cluster membership. (b). MDS based on the –log10p(c-scores) matrix of pathways colored by cluster membership. (c). Co-membership heatmaps of significantly concordant pathways within each pathway cluster between each pair of studies. Each cell is colored by the proportion of pathways in which the two corresponding studies are clustered together based on the pathway level c-scores. Each heatmap is annotated with key noun phrases from the text mining algorithm.

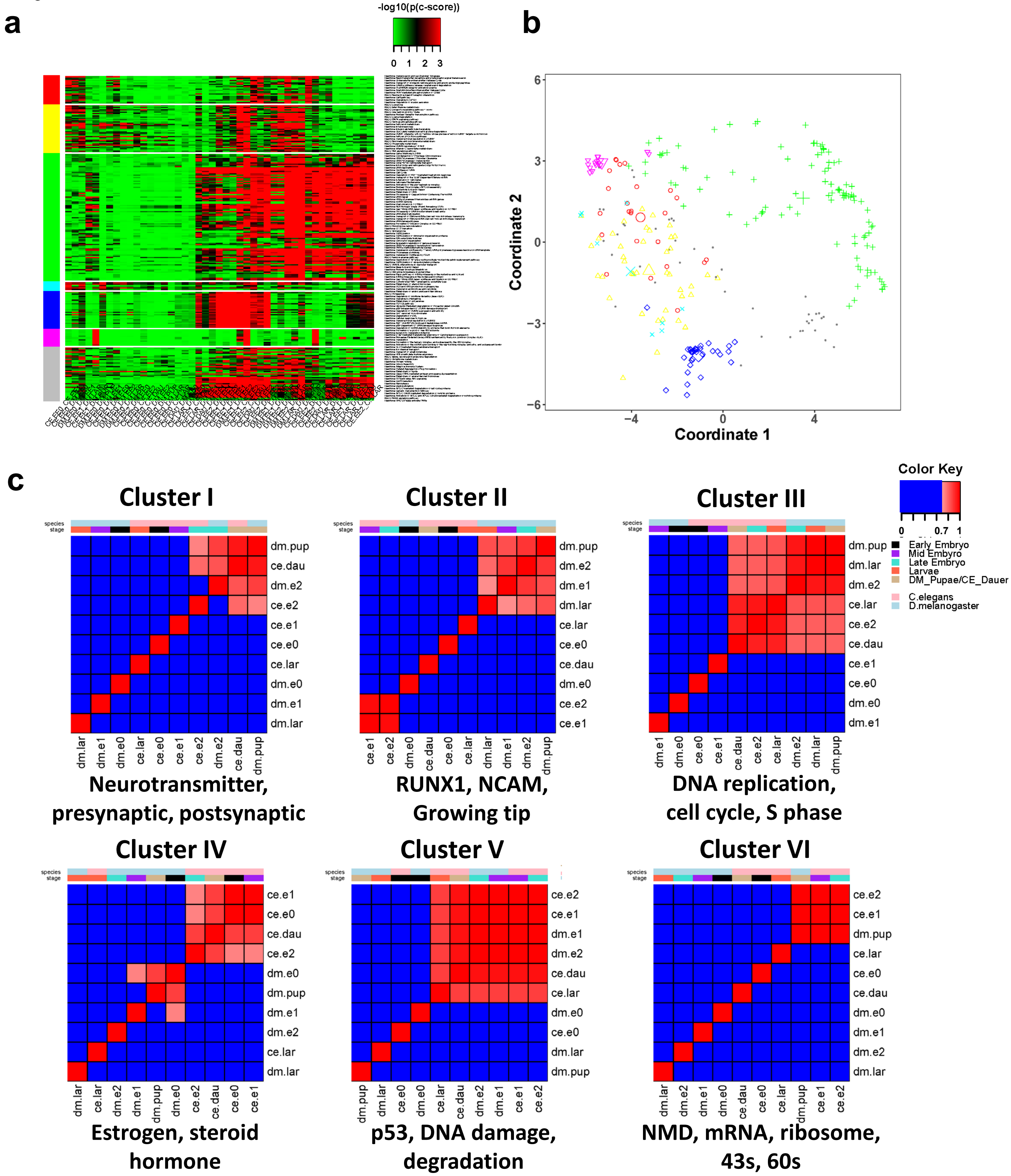

**Supplementary Figure 8.** Results from Case Study 2. (a). Summary of DE evidence with the pathway level c-scores (orange in the upper right region) and d-scores (blue in lower left region). X and y axes represent the average DE posterior probabilities, size of dots represent the magnitudes of c-scores (orange) or d-scores (blue). ce.e2: Late embryo stage of CE; ce.dau: Dauer stage of CE; dm.e2: Late embryo stage of DM; dm.pup: Pupae stage of DM. Three example pathways are highlighted using different shapes ("△": cel03440 – KEGG: Homologous recombination; "◇": cel03430 – KEGG: DNA mismatch repair; "□": R-CEL-168643 - Reactome: Nucleotide-binding domain, leucine rich repeat containing receptor (NLR) signaling pathways). Orange represents concordant pathways, while blue represents discordant pathways. (b). Gene-wise heatmap of posterior mean of DE indicators of the ce.e2-dm.e2 in cel03440, ce.e2-dm.e2 in cel03430 and ce.dau-dm.pup in R-CEL-168643.

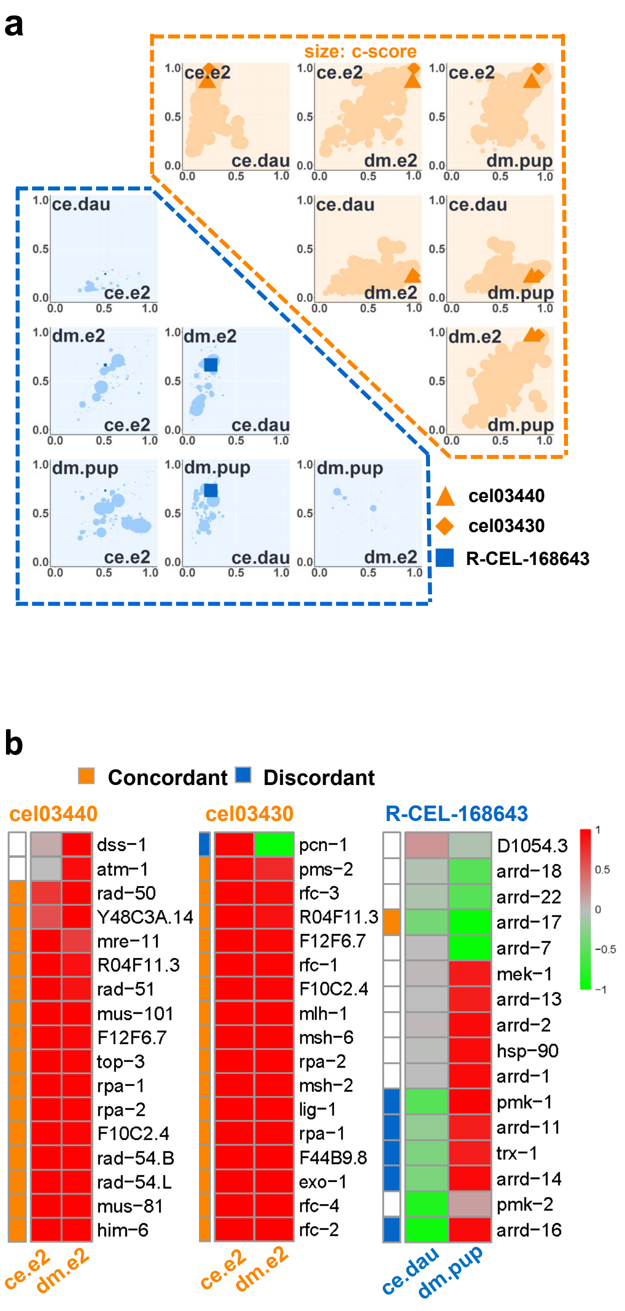

**Supplementary Figure 9.** Pathway topology plots of the selective pathways Case Study 2: (a). cel03440 (ce.e2-dm.e2), (b). cel03440 (ce.e2-dm.e2) and (c). R-CEL-168643 (ce.dau-dm.pup). Due to the complexity in Reactome pathways, not all genes are spelled out in the topological annotation. We highlight all the involved genes in orange parentheses.

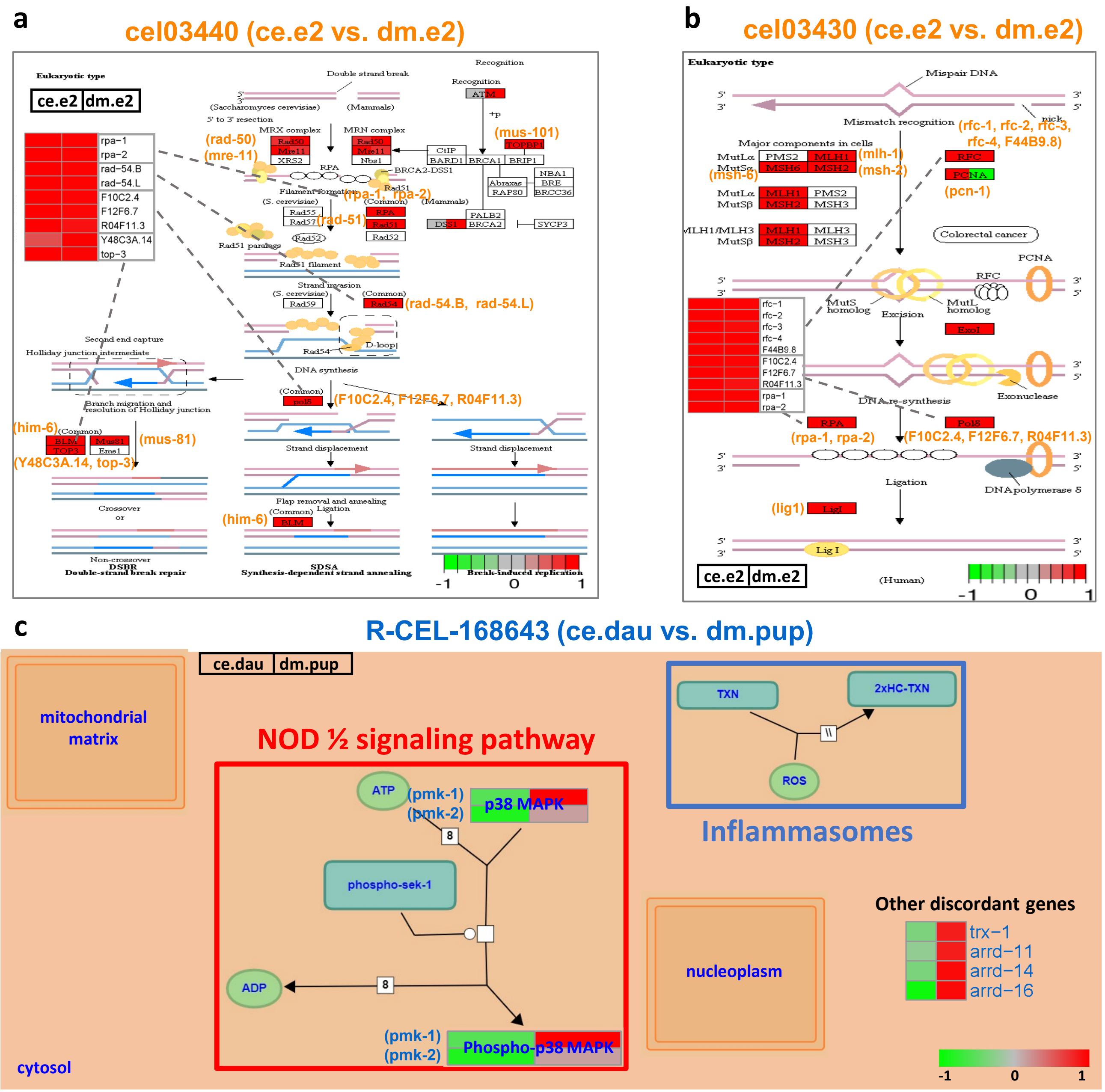

**Supplementary Figure 10.** The p-value elbow plots from the local discordant community detection algorithm for pathway examples hsa04670 (HS-MS) and hsa04662 (HS-MS) Case Study 1. In each plot, the red dashed line denotes the –log10 transformation of the minimum p-value minus its 2 standard deviation (module size is 4 in hsa04662 and 11 in hsa04670). The largest module size whose –log10(p-value) above this dashed line is selected (annotated by red triangles; 6 in hsa04662 and 12 in hsa04670).

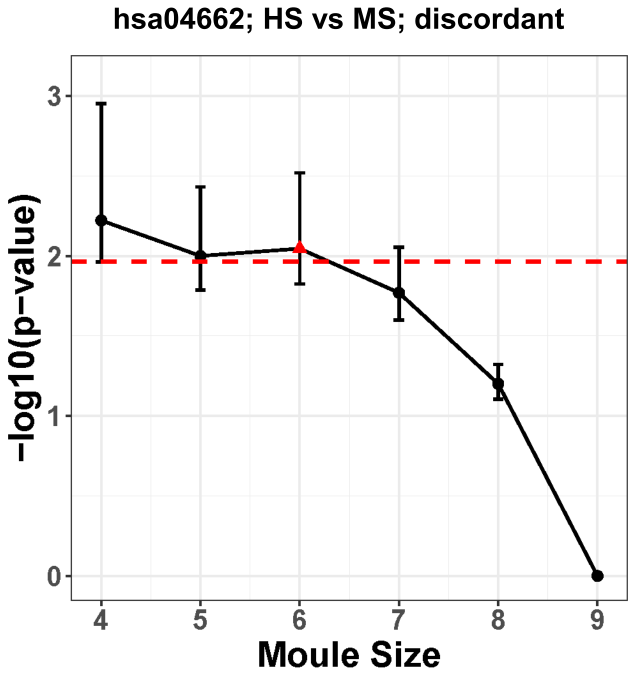


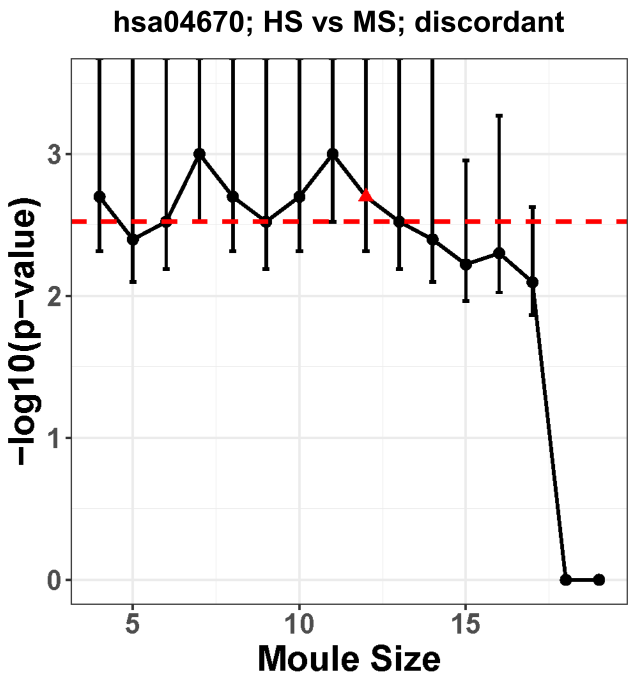

**Supplementary Table 1:** 12 inflammatory response studies in human and mouse. Numbers in each parenthesis denote the disease sample size versus the control sample size in the corresponding dataset.

| | **human** | **mouse** |
|---|---|---|
| Burn | GSE37069 (553 vs 37) | GSE7404 (16 vs 16) |
| Infection | GSE30119 (99 vs 44) | GSE20524 (55 vs 17) |
| Trauma | GSE36809 (820 vs 37) | GSE7404 (16 vs 16) |
| Sepsis | GSE13904 (106 vs 18) | GSE19668 (25 vs 25) |
| Endotoxin (LPS) | GSE3284 (66 vs 44) | GSE7404 (8 vs 8) |
| Acute Respiratory Distress Syndrome (ARDS) | GSE10474 (13 vs 21) | GSE52684 (17 vs 13) |

**Supplementary Table 2:** Differences of analysis strategies and subjective thresholds in the two contradictory PNAS papers. DE genes from human only or intersection of human & mouse are taken with certain DE selection criteria to calculate Pearson or Spearman correlations. Their data sources are also different. Data and programming code are not available in both papers to replicate the findings.

| | **Seok et al. (2013)** | **Takao et al. (2014)** |
|---|---|---|
| DE genes of interest | Human only | Intersection of human & mouse |
| DE selection criteria | FDR < 0.001, FC≥2 | p-value<0.0001, FC≥2 (H)<br>p-value<0.0001, FC≥1.2 (M) |
| Correlation measure | $R^2$ (squared Pearson) | ρ (Spearman) |
| Data source | First-hand data from experimental labs | Outputs (p-value, FC) from NextBio |
| Data and programming code | Not available | Not available |

**Supplementary Table 3**. See “Supplementary Table 3.xlsx”. Genome-wide c-scores and d-scores (with associated p-values) for all pairs of 12 human and mouse studies in Case Study 1. In the c-score table, HB, HI, HT and HS resembled each other well with genome-wide pair-wise c-scores ranging from 0.25 to 0.52 (highlighted in dark green). HA and HL has almost no resemblance with c-scores≈0 (highlighted in light green). MI, MB and MT are overall more similar to the four human studies (HB, HI, HT and HS) in a weaker congruence level with c-scores=0.081-0.2 (highlighted in pastel pink). Within each inflammatory disease, only MI-HI and MB-HB have relatively large c-scores at 0.2 and 0.11 (red), while c-scores of MS-HS, MT-HT, MA-HA and ML-HL are almost 0 (blue). The six mouse studies generally do not mimic each other (highlighted in light blue).

**Supplementary Table 4**. See “Supplementary Table 4.xlsx”. Text mining results of pathway clusters identified in Case Study 1.

**Supplementary Table 5**. See “Supplementary Table 5.xlsx”. Genome-wide c-scores and d-scores (with associated p-values) for all pairs of 10 studies in Case Study 2. In the c-score table, ce.e0-dm.e0, ce.e1-dm.e1, ce.e2-dm.e1, ce.e2-dm.e2 show within-stage concordance (highlighted in yellow; c-scores≥0.1) and ce.lar-dm.lar has slightly weaker within-stage concordance (highlighted in green; c-scores=0.087) while ce.e2- dm.lar, ce.e2-dm.pup and ce.dau-dm.e2 show cross-stage concordance (highlighted in purple; c-scores≥0.1).

**Supplementary Table 6**. See “Supplementary Table 6.xlsx”. Text mining results of pathway clusters identified in Case Study 2.

**Supplementary Table 7**. Confusion matrix of DE gene status comparing between a human study (H) and a mouse study (M). $\Omega^{H+}$, $\Omega^{H-}$ and $\Omega^{H0}$ are collections of up-regulated, down-regulated and non-differential genes in human, where $\Omega^{H+} = \{g: \delta_g^H = 1\}$, $\Omega^{H-} = \{g: \delta_g^H = -1\}$ and $\Omega^{H0} = \{g: \delta_g^H = 0\}$, and similarly for mouse. Denote by $a$, $e$ and $i$ the number of cross-species DE concordant genes: $a = \#(\Omega^{H+} \cap \Omega^{M+})$ (number of concordant up-regulated genes), $e = \#(\Omega^{H0} \cap \Omega^{M0})$ (number of concordant no-change genes), and $i = \#(\Omega^{H-} \cap \Omega^{M-})$ (number of concordant down-regulated genes). $b, c, d, f, g, h$ are defined similarly.

| | $\Omega^{H+}$ | $\Omega^{H0}$ | $\Omega^{H-}$ | sum |
|---|---|---|---|---|
| $\Omega^{M+}$ | a | b | c | A |
| $\Omega^{M0}$ | d | e | f | B |
| $\Omega^{M-}$ | g | h | i | C |
| sum | D | E | F | G |

**Supplementary Note 1.** SA algorithm in the local module detection given an initial module.

**Data:** Initial module $G_m^c$, $S_k$, $G$, $T_0 = 10$, $T_f = 1e - 5$, $\mu = 0.95$, $N = 1000$, $R = 10000$

**Result:** A concordant/discordant gene set of size $m$ with an optimized average shortest path value.

**while** $run < R$ *&* $count < N$ *&* $T_0 > T_f$ **do**

- $run \leftarrow run + 1$;
- Propose a new gene set by randomly substituting one gene in $G_m^c$ to a gene randomly drawn from the full concordant/discordant gene set $G$ and calculate its $avgSP(G_m^t)$ ;
- **if** $avgSP(G_m^t) < avgSP(G_m^c)$ ***or*** $avgSP(G_m^c) = \infty$ **then**
  - $G_m^c \leftarrow G_m^t$;
  - $avgSP(G_m^c) \leftarrow avgSP(G_m^t)$;
- **else**
  - $count \leftarrow count + 1$;
  - Acceptance probability $r = \min(1, \exp\{\frac{-[avgSP(G_m^t) - avgSP(G_m^c)]}{T_0}\})$ ;
  - Draw a random number $u \sim Unif(0, 1)$;
  - **if** $u > r$ **then**
    - $T_0 \leftarrow T_0 \times \mu$;
  - **else**
    - $G_m^c \leftarrow G_m^t$;
    - $avgSP(G_m^c) \leftarrow avgSP(G_m^t)$;
  - **end**
- **end**

**end**

**Supplementary Note 2.** Overall SA algorithm in local module detection.

**Data:** $S_k$, $G$, $T_0 = 10$, $T_f = 1e-5$, $\mu = 0.95$, $N = 1000$, $R = 10000$, $x = 100$, $y = 10$
**Result:** Top $y$ concordant/discordant gene sets of size $m$ with the smallest average shortest path values among the $x$ repetitions.
**if** *Top* $y$ *concordant/discordant gene sets of size* $m-1$ *are provided as* $\{G^i_{m-1}\}_{i=1:y}$
**then**
  **for** $i$ = *1, 2,...,*$y$ **do**
    **for** $j$ = *1, 2,...,*$x/y$ **do**
      Set the initial module as $\{G^i_{m-1}, g_j\}$ where $g_j$ is a gene randomly selected from $G$. Run Algorithm 1 with this initial gene module and record the output gene module as $G^{ij}_m$
    **endfor**
  **endfor**
  The top $y$ gene modules with the smallest average shortest path values among $\{G^{ij}_m; 1 \leq i \leq y, 1 \leq j \leq x/y\}$ are stored and passed to next $m+1$ scenario. The best of them is output as the final result at module size $m$.
**else**
  **for** $i$ = *1,...,*$x$ **do**
    Randomly draw $m$ genes from $G$ as an initial module and run Algorithm 1 with this initial module. The result is recorded as $G^i_m$
  **endfor**
  The top $y$ gene modules with the smallest average shortest path values among $\{G^i_m; 1 \leq i \leq x\}$ are stored and passed to next $m+1$ scenario. The best of them is output as the final result at module size $m$.
**endif**
